# Focal vs Diffuse: Mechanisms of attention mediated performance enhancement in a hierarchical model of the visual system

**DOI:** 10.1101/2023.05.09.540022

**Authors:** Xiang Wang, Monika P. Jadi

**Affiliations:** Interdepartmental Neuroscience Program, Yale University, New Haven, CT 06511; Department of Psychiatry, Yale University, New Haven, CT 06511; Department of Neuroscience, Yale University, New Haven, CT 06511

## Abstract

Spatial attention is an essential cognitive process for visual perception, especially in complex scenes with poor luminance. Attentional modulation of neural activity has been documented across the visual cortex. However, how these changes in neural response lead to better behavioral performance on challenging tasks remain unknown. In our study, we implemented spatial attention in a deep convolutional neural network model of the ventral visual hierarchy and measured its impact on categorization performance on cluttered images with varying contrast levels. We applied attention to network units in three ways: enhancement in attended region (EAR), suppression in unattended region (SUAR) or both. When focally applied to a single convolutional layer, SUAR is more effective in boosting performance than EAR, especially in the presence of high contrast distractors. SUAR is also effective in recovering degraded performance due to low contrast targets, whereas EAR fails to recover. These results predict a novel mechanism of suppression of neural activity corresponding to the unattended parts of visual space. Intriguingly, EAR in our model achieves the same performance as SUAR when attention is diffusely applied to stacks of successive convolutional layers, irrespective of target contrast. This suggests an alternate mechanism of attention wherein enhancement of attended neural activity alone, when applied to successive cortical encoding stages, is an effective strategy for boosting performance in challenging object recognition tasks. Our results indicate that the two alternative attentional mechanisms are functionally equivalent in tackling challenging object recognition tasks.

## INTRODUCTION

Our visual system is bombarded with many stimuli every moment. Selecting which stimulus should be prioritized for processing depends on the task at hand and requires visuo-spatial attention.^1^ Experiments aiming to explore the neural correlates of attention have characterized various attentional modulations of neural activity in visual cortex. For example, spatial attention modulates the variability of spiking, making neurons more reliable across trials and less correlated with others.^2, 3^ Within the framework of signal processing, such modulation is interpreted as reducing the “local” noise that is present in the neural signal representing the attended region. Attention could also act in a push-pull manner in visual space: enhance neural responses within the attended region^4–6^ and/or suppress those in the unattended region.^7–12^ The suppressive effect could reduce the “global” noise that represents the clutter in the visual scene,^13^ solving the routing problem^14^ that the signal traversing the hierarchical network may get contaminated by surrounding noise because of the convergence of information inherent in the network. However, how this push-pull mechanism is deployed in cortical space across stages of the visual hierarchy remains unclear. Although the electrophysiological literature documents attentional enhancement of visual activity across multiple brain areas,^6, 15–18^ such studies pertaining to visual suppression are lacking, with existing ones mainly focusing on the striate cortex.^8, 9^ Functionally, since the consequences of enhancing and suppressive computations can be sensitive to where they are applied in a hierarchy,^19–22^ it is not known whether and how the push-pull mechanism of attention exerted in a hierarchical system could facilitate its behavioral performance. Empirically, it is challenging to systematically characterize attention mechanisms across the visual and cortical spaces. Computational models can therefore be a powerful approach for investigating these questions.

Deep convolutional neural networks (DCNNs), inspired by the computations and the hierarchical architecture of the visual system, have become start-of-the-art models of human vision.^23–25^ At the behavioral level, feedforward DCNNs achieve human-level categorization performance after training on large volumes of hand-annotated images (ImageNet).^26, 27^ This allows a direct comparison between model performance and that of primates for testing the effects of attention, a task that is not possible for other models of attention. At the neuronal level, these task-optimized DCNNs feature sensory representations predictive of neural representations along the primate ventral stream pathway.^28–33^ Adding recurrent and feedback mechanisms to the canonical feedforward architecture of DCNNs further revealed the computational roles of inter-area and within-area connections.^34–38^ Comparing DCNNs and the visual system processing also generated insightful theories about neural mechanisms behind visual computations such as scene segmentation,^39^ shape/category encoding,^40^ and emotion selectivity.^41^ Given their success in modeling the primate ventral stream, DCNNs have the potential to uncover the computational principles that lead to perceptual improvement mediated by attention. Indeed, recent work showed that implementing the neural computations found in feature attention^18, 42^ in these networks was effective at improving performance on more challenging visual tasks such as classifying overlapping or cluttered images^19^.

In this study, we investigated the functional consequences of the push-pull mechanism of spatial attention deployed in visual and cortical space by characterizing its influence in a DCNN model. We applied enhancement of neural activity representing the attended region (EAR) and suppression of the same for the unattended region (SUAR) to different parts of the model hierarchy and evaluated its categorization performance on cluttered images with varying contrast levels. When attention was applied to a single layer of the DCNN, we found that SUAR boosted the object recognition performance to the same level as the one on single images, while EAR failed to recover the degraded performance. On the other hand, when multiple successive layers were modulated by attention, the contributions of EAR and SUAR were undifferentiated; either one was sufficient to recover the performance. Performance benefits induced by the two mechanisms aligned with their impacts on the information transmission efficiency of the network. The combination of the two mechanisms provide a novel perspective for explaining the large improvements in psychophysical performance by modest neuronal gain changes due to attention. While EAR has been extensively investigated in neurophysiological studies, our results identify SUAR and multi-stage EAR mechanisms as targets for future studies of visual attention.

## RESULTS

### Performance of DCNNs is sensitive to image contrast

We use VGG-16^19, 43^, a DCNN variant, to study the effect of neurally identified attention mechanisms on object recognition performance. The architecture consists of 5 “stacks” of convolutional and max pooling layers, followed by 2 fully connected layers and a series of binary classifiers (Figure 1A, Methods). VGG-16 was pre-trained on standard ImageNet images, and has been shown to exhibit near-human-level performance on object categorization tasks and with feature representation that corresponds closely to single-voxel activity of neurons across the ventral stream.^30^ We used a series of binary classifiers (20 in total) instead of the originally proposed softmax layer in VGG-16 to speed up the evaluation process. The weights of binary classifiers were learned by training on the single-category images from the ImageNet validation set (Methods).^19^

**Figure 1.**
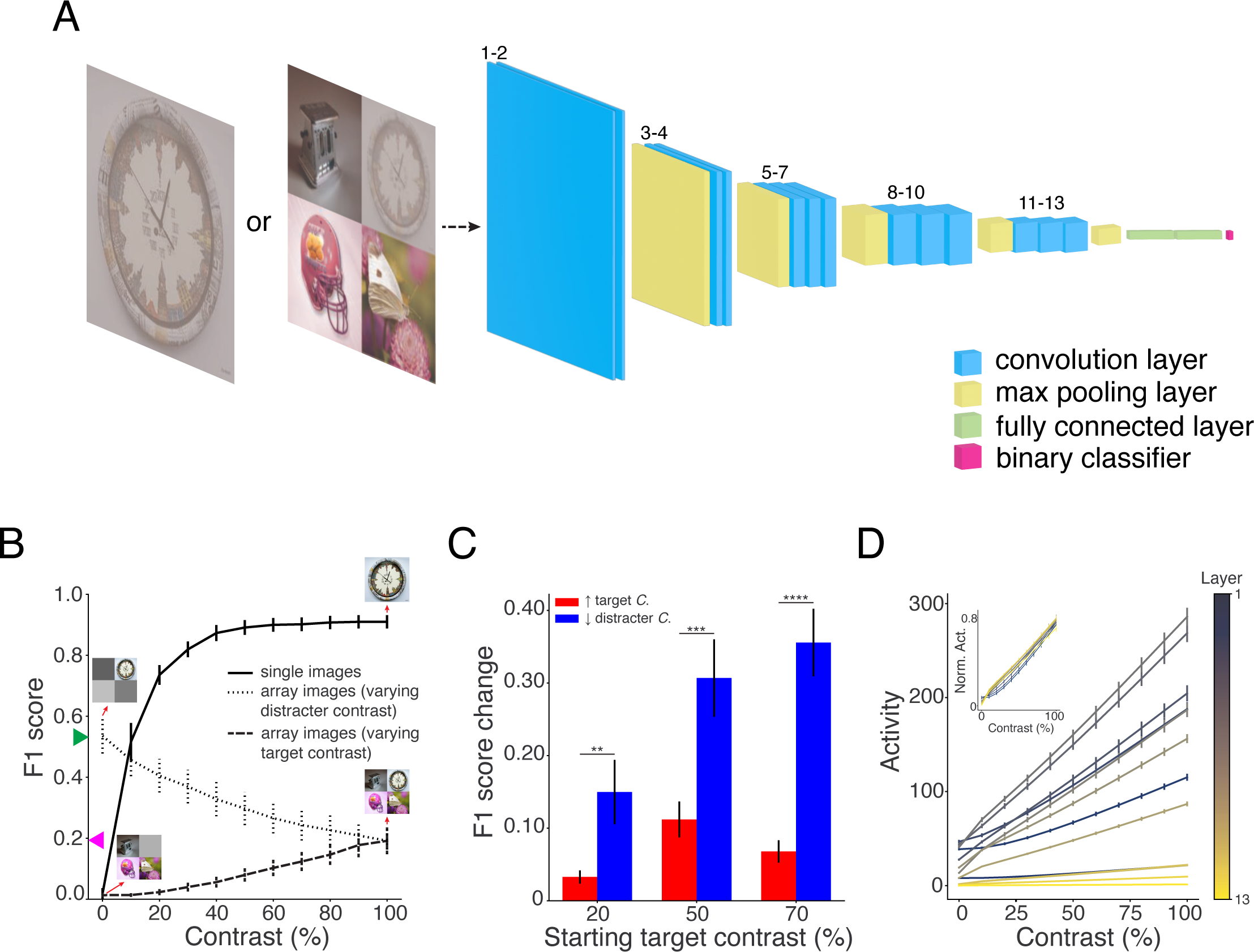
Network structure and the impact of image contrast on model performance. (A) We fed low-contrast images (single-category image or composite image) to a pre-trained deep convolutional neural network, VGG-16. The network consists of 13 convolutional layers (labeled in numbers), 5 max pooling layers, and 2 fully connected layers. The weights of the network were obtained by optimizing the model’s performance on the 1000-way classification task of the ImageNet dataset.^43^ The last layer of the network is a series of binary classifiers (logistic regression), one for each of the 20 testing categories. These classifiers were trained on the ImageNet validation set. (B) Network performance on single-category images or composite images with different contrasts. We modified the contrast level of either single-category images (solid line), or distracters in composite images (dotted line), or targets in composite images (dashed line) and measured the network classification performance on 20 categories (Mean ± SEM). (C) Performance change (Mean ± SEM) induced by increase in target contrast (red) or decrease in distracter contrast (blue). From a composite image set with low-contrast targets (20%, 50%, or 70%) and full-contrast distracters, we either boosted the target contrast by 100% (no more than full contrast) or reduced the distracter contrast by 100% (down to 0% of contrast) and quantified the performance change of the network. The number of asterisks denotes the p-value cutoff of the difference between the two contrast manipulations (Wilcoxon signed-rank test, *N* = 20), where * = *p* < 0.05, ** = *p* < 0.01, *** = *p* < 0.001, and **** = *p* < 0.0001. See more illustration and detailed results in Figure 1—figure supplement 1. (D) Neural activity of convolutional layers (normalized in the inset) in response to single-category images with different contrasts. The reported neural activity was the mean over spatial dimensions, feature maps, and 20 categories (Mean ± SEM).

To determine if image quality could control the difficulty of the object categorization task, we input ImageNet validation images with modified contrast levels into the network (Figure 1A, Methods). The model yielded saturated performance at 40% contrast, and a steep decrease in performance with further loss in image contrast (Figure 1B, solid line). This function approximated the behavioral performance curve (psychometric function) in primates,^44–46^ which indicated the feasibility of manipulating contrast to control task difficulty. In order to apply spatial attention to the model, we next created a relatively “crowded” scene by scaling down ImageNet images and arranging a target-category image with 3 distracters from other categories on a two-by-two grid (composite images).^19^ Given the information loss brought by size scaling, the network’s performance dropped to 58.8% of its peak when distracter images had 0% contrast (Figure 1B, leftmost point of the dotted line). Increasing the contrast level of distracters further degraded the model’s performance (Figure 1B, dotted line). The performance was also a function of the target contrast (Figure 1B, dashed line); however, this showed little saturation, given the significantly reduced peak performance for composite images.

The contrast manipulations on composite images (Figure 1B) demonstrated that both the target contrast and the distracter contrast influenced the classification accuracy of the DCNN. The network appeared to be more sensitive to the contrast change of distracters than that of the target. A contrast change of 100% in distracters led to a performance difference of 0.34 while the same contrast change in the target altered the performance by 0.18. To further quantify this observation, we measured the model’s performance difference in response to an increase in the target contrast or a decrease in the distracter contrast from the same composite image (Figure 1—figure supplement 1A). Consistent across different initial conditions, both manipulations improved the model’s performance, but their magnitudes of improvement differed when the contrast change was substantial (> 90%, Figure 1—figure supplement 1B). When changing the contrast by 100%, decreasing the distracter contrast always outperformed increasing the target contrast, regardless of the initial contrast difference between them (Figure 1C). Therefore, under the task of classifying composite images, the neural network was more sensitive to the distracter content than the target content.

In non-human primates, the neural activity in early and mid-tier visual processing stages, such as V1 and V4, increases monotonically with the stimulus contrast.^5, 47^ To see if that pattern holds for DCNNs, we recorded the mean activity of artificial units in every convolutional layer in response to ImageNet images of varying contrasts. Notwithstanding the difference in absolute activity value across layers, the mean activity of convolutional units increased linearly with the image contrast for every layer (Figure 1D). Attention-mediated response gain for low-contrast stimuli is believed to elevate the signal-to-noise ratio of neurons that encode the attended stimulus and, therefore, compensate for the information loss caused by the contrast reduction.^3, 48^ We thus hypothesized that the firing-rate modulatory effects of spatial attention observed in neurophysiology would also benefit the classification performance of the artificial neural network on images with different contrast levels.

### DCNN performance is differentially modulated by components of spatial attention mechanism

We simulated spatial attention in VGG-16 as “attending” to the target category region of a composite image. We applied the attentional modulatory effect to an individual convolutional layer by scaling the slopes of the activation functions of all feature maps (Figure 2A). In line with empirical observations in the visual cortex^18, 49–51^, we applied multiplicative scaling to the activation functions (Methods). We first implemented a scenario with attentional modulation of the entire visual space (EAR+SUAR), in which the neural activity was enhanced for artificial units within the corresponding attended quadrant (EAR) and suppressed for those within the other 3 unattended quadrants (SUAR). To determine the most effective layer for spatial attention, we evaluated the network’s performance on full-contrast composite images when attention was applied to different network layers. For every condition, we tested a range of attention strengths (absolute value of the multiplicative factor) and selected the highest performance (Figure 2B, Figure 2—figure supplement 1A). Spatial attention improved the network performance only when the target category was “attended” (Figure 2—figure supplement 1A). The greatest improvement occurred when the last convolutional layer was modulated (0.53 increase in F1 score, Figure 3A), which was consistent with a previous study implementing feature attention in the DCNN.^19^ Simultaneously modulating all layers led to a slightly better performance than modulating the last layer only (0.023 increase in F1 score, Figure 2—figure supplement 1C), but it was not statistically significant (Wilcoxon signed-rank test, *p* > 0.05, *N* = 20). Thus, we limited further analyses to attention implementation at individual layers for better interpretability.

**Figure 2.**
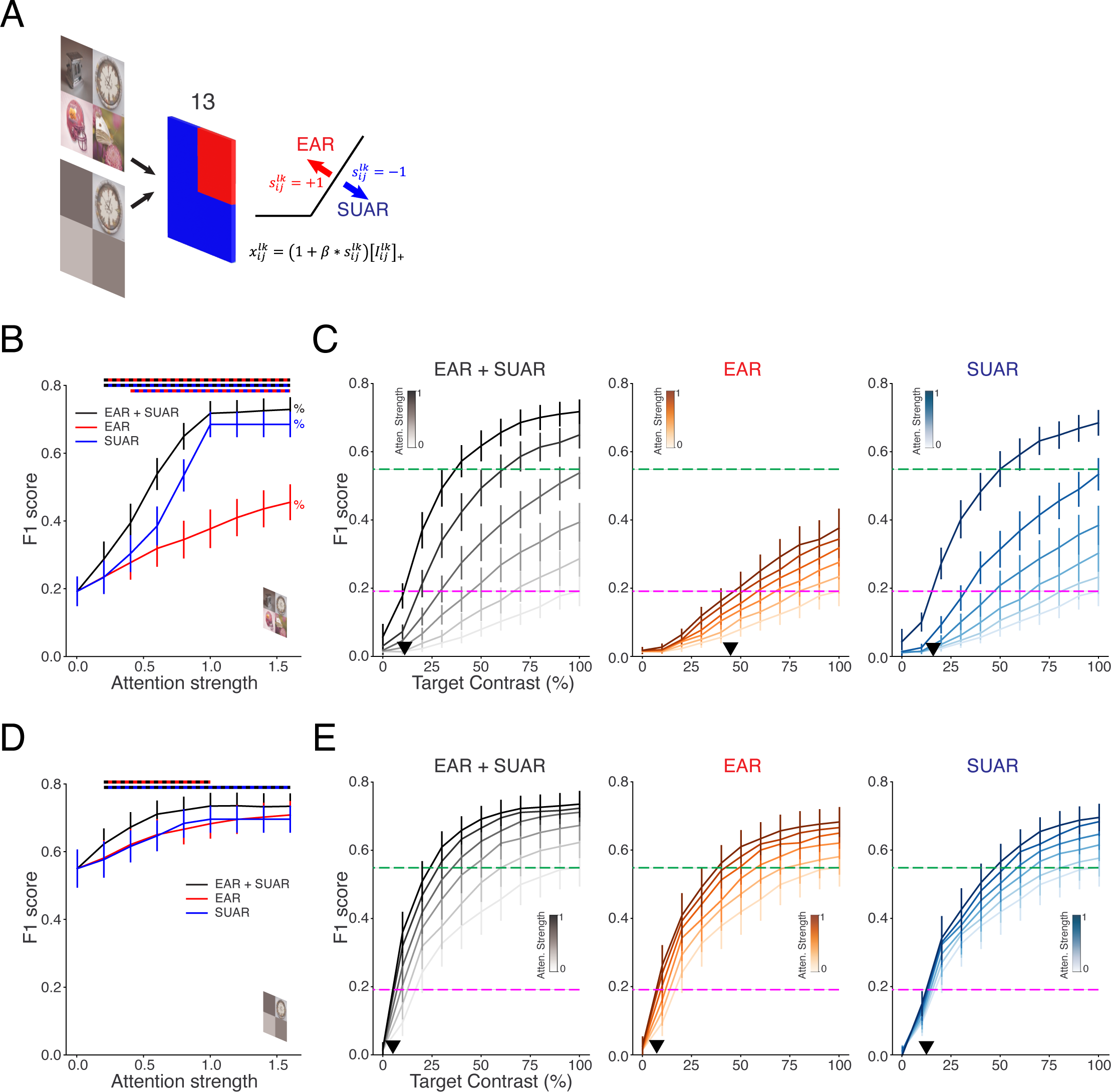
Effect of focal vs. diffuse attention applied to individual layer of network. (A) Schematic of the two attention effects that could be applied to a convolutional layer: For all units across feature maps, the slopes of their activation functions were altered based on an overall strength value β. The direction of modulation was decided by whether they resided in the attended region 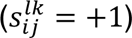 or the unattended region 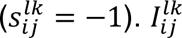 was the input from the previous layer. We applied attention effects to the last convolutional layer (layer 13) of VGG-16 since it achieved the highest performance (Figure 3A). The distracter contrast of the input was either 100% (B, C) or 0% (D, E). (B) Network performance on full-contrast composite images when spatial attention of different strengths was applied to layer 13. Attentional modulation was either EAR+SUAR, EAR, or SUAR. The reported performance was averaged across 20 categories (Mean ± SEM). Performances at each attention strength were compared among different attention conditions and the significance was indicated by the dual-color dotted line on top (black-red: EAR+SUAR vs. EAR; black-blue: EAR+SUAR vs. SUAR; red-blue: EAR vs. SUAR; Wilcoxon signed-rank test, *N* = 20, Bonferroni corrected, *n* = 3, *p* < 0.0167). % indicates a significant difference in performance among different attention strengths for a specific attention condition (Friedman test with Conover post hoc pairwise comparisons, *N* = 20, Bonferroni corrected, *n* = 9, *p* < 0.00555 for any pair of comparison). (C) Attentional improvement of the network performance on composite images with different target contrast. Attention was implemented with varying strengths (a step of 0.2) and different modulatory types (EAR+SUAR, EAR, SUAR). The performance references without attention are shown: 100% contrast of target and 0% contrast of distracters (green dotted line); 100% contrast of target and distracters (magenta dotted line). Black triangle marks the lowest contrast level of the target (maximal information loss) that could be recovered by attention, which is the contrast value of the intersection between the curve for the highest attention and the magenta dotted line. (D) and (E) Same as (B) and (C) respectively, but on control images where the contrast of distracters was set to 0%.

**Figure 3.**
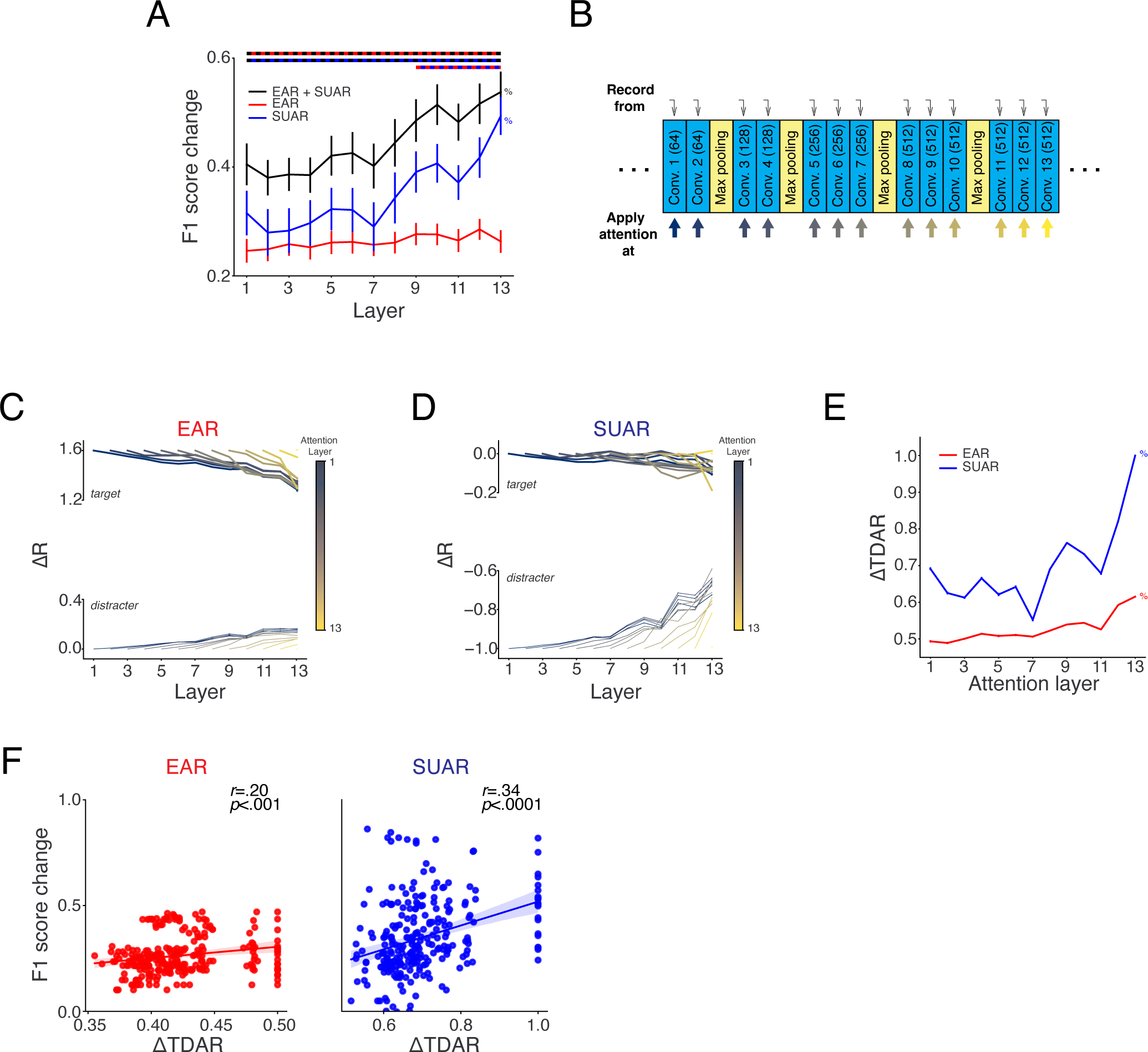
Attention-induced activity modulation along the network hierarchy. (A) Average change (Mean ± SEM) in binary classification performance as a function of layer to which spatial attention (EAR+SUAR, EAR, SUAR) was applied. For all cases, we used the best performing strength from the range tested. Horizontal bars on top indicate layer numbers for which there was a significant difference, with color code indicating the pairs of modulatory types considered for comparison (Wilcoxon signed-rank test, *N* = 20, Bonferroni corrected, *n* = 3, *p* < 0.0167). % indicates a significant difference in performance among different layers for a given modulatory type (Friedman test with Conover post hoc pairwise comparisons, *N* = 20, Bonferroni corrected, *n* = 13, *p* < 0.00384 for any pair of comparison). (B) Recording setup. The spatially averaged activity of each convolutional layer was recorded while attention was applied to every layers individually. Activity was in response to full-contrast composite images. (C) Relative activity change (ΔR, Mean ± SEM, error bar is too small to be seen) in the target quadrant (thick lines) or distracter quadrants (thin lines) of each layer when EAR was applied to individual layers. Only the layers following the affected location are shown. The activity was averaged over spatial dimensions and feature maps for a given category. The activity change was computed per category and reported as the average across categories. (D) Same as (C) for SUAR. (E) Relative change (Mean ± SEM, error bar is too small to be seen) of target-distracter activity ratio (ΔTDAR) derived from the neural activity change (ΔR) in layer 13 when EAR (red) or SUAR (blue) was applied to a layer. The TDAR is defined as the negative ratio of the distracter region activity over the target region activity (see Methods for details). % indicates a significant difference in ΔTDAR among layers (Friedman test with Conover post hoc pairwise comparisons, *N* = 20, Bonferroni corrected, *n* = 13, *p* < 0.00384 for any pair of comparison). Significant differences in ΔTDAR were detected between SUAR and EAR for every layer (Wilcoxon signed-rank test, *N* = 20, *p* < 0.05). (F) Relationship between performance change and ΔTDAR of layer 13, for EAR or SUAR. Every data point is an instance for a category with attention applied to an individual layer.

To determine whether the behavioral benefit of spatial attention depends on image contrast, we applied attention effects to the last convolutional layer (since it manifested the greatest attentional benefit) and input composite images with different contrast levels of the target (Figure 2A). We found that the model’s accuracy was boosted at every contrast level (Figure 2C, left panel). Attention could compensate for the performance loss from the image’s poor quality, boosting the accuracy for contrast level as low as 12% to the level for full-contrast images (Figure 2C, magenta dotted line). The boosted performance could even surpass the network’s baseline performance on composite images with minimal clutter (Figure 2C, green dotted line). These results demonstrated the effectiveness of spatial attention at counteracting the signal degradation from poor image quality (low contrast) and reducing the influence of “global” noise (distracters) that causes cluttering.

The spatial attention we implemented above included enhancement of neural activity in the attended region (EAR) and suppression in unattended regions (SUAR), whose relative contributions to visual perception remain unknown. Noting that the activity of the network was linearly dependent on image contrast (Figure 1D), and that the network benefited more from the contrast reduction of distracters than the contrast enhancement of the target (Figure 1C), we hypothesized that the two types of attentional modulation may produce heterogenous behavioral outcomes, and that SUAR might lead to a better performance than EAR.

To test our hypothesis in the context of DCNNs, we examined the behavioral influence of SUAR and EAR by applying the two effects separately on a convolutional layer of the network. For a range of attention strengths, SUAR always outperformed EAR (Figure 2B, Figure 2—figure supplement 1A). SUAR could also lead to a better performance on low-contrast images than EAR, as reflected by a higher peak performance and a lower contrast level that could be fully recovered (Figure 2C, middle and right panels). These results demonstrated that distracter suppression provides more behavioral benefits than target enhancement for artificial neural networks categorizing objects in cluttered scenes. The clutter level (the amount of “global” noise) should influence the difference in performance brought by distracter suppression. We tested the network on a control image set where the clutter was minimized by zeroing out the contrast of distracters. As expected, the benefit from SUAR was much lower on the control dataset, regardless of the attention strength or the target contrast level (Figure 2D, E). In fact, the impact of SUAR was highly dependent on the clutter level (as measured by the distracter contrast in Figure 2—figure supplement 1D), while EAR was not. Moreover, every layer showed negligible performance improvement on control images by applying suppression only (Figure 2—figure supplement 1B). These observations suggested that SUAR reduces the interference of distracters, thereby improving the network’s performance under the setting of an effective crowded scene.

### Attentional components show distinct effects on signal transmission

We observed that the influence of EAR+SUAR attention on performance varied with the network layer of its implementation (Figure 3A). Also, distracter suppression played a major role in attentional improvement (Figure 2B, 2C). Thus, we were interested in how the two types of attentional modulation contribute to the performance when applied to different locations. To answer this, we repeated the layer-wise performance analysis for EAR and SUAR separately (Figure 3A). Performance improvement from EAR was lower than the EAR+SUAR attention and remained the same across layers, but the benefit of SUAR was greater for deeper layers. Although it was still lower than the performance of EAR+SUAR attention, the trend of SUAR was similar to that of EAR+SUAR attention, again substantiating our conclusion that distracter suppression was the major contributor in the attentional facilitation of behavior (Figure 2B).

To understand the computational mechanism that led to the behavioral difference between EAR and SUAR, we explored how the two strategies affect the neural activity of the convolutional layers. We applied EAR or SUAR with optimized attention strength to each convolutional layer and “recorded” relative activity changes (ΔR) in the target region (signal) or the distracter region (noise) of each layer (Figure 3B). As the modulatory effect of EAR propagated through the network, the activity enhancement in the target region gradually decayed while the activity in the distracter region increased (Figure 3C). This was because of the extended spatial sensitivity in the deeper layers due to the convolution and max pooling operations. As a result of the same reason, the activity change in the distracter region from SUAR also faded as it propagated to the deeper layers (Figure 3D). To quantify their effects on the network’s information transmission, we assumed that the target signal was mostly encoded in the target region activity while the distracter (noise) signal was mostly encoded by the distracter region. Therefore, the signal-to-noise ratio of the network can be approximated by the target-distracter activity ratio (TDAR, see Methods for the formula). We computed the relative change in the target-distracter activity ratio (ΔTDAR) of layer 13 from ΔR of the target or distracter region (Figure 3C, D). We focused on the activity of this last convolutional layer because it’s the closest one to the classifiers and outputs the filtered information from all convolutional layers. We found that EAR exhibited modest increase in TDAR when applied to “late” layers. However, SUAR applied to “late” layers greatly improved the ratio compared to its application to “early” layers (Figure 3E). Interestingly, ΔTDAR for SUAR was always higher than that for EAR, which agreed with the performance difference between the two strategies (Figure 3A). In addition, the layer dependence of TDAR had a similar pattern to that of performance (Figure 3A). A significant positive correlation was found between performance change and ΔTDAR for both strategies (Figure 3F). Such correlations validated our metric as an appropriate estimate of the model’s signal quality during information transmission. Collectively, these results suggested that EAR and SUAR provided distinct advantages for the model’s signal processing. The superior effectiveness of SUAR and its layer dependency may emerge from the more informative signal transmission triggered by its noise reduction.

### Attentional modulations on multiple consecutive layers show distinct properties

The location to which we applied attentional modulation thus far was limited to individual convolutional layers. Since attention modulatory effects have been observed in multiple brain areas in the primate visual system, ^6, 15–18, 52, 53^ we next investigated the functional consequences of implementing attentional modulation to multiple convolutional layers in the DCNN. Previous work mapping visual representations in the primate visual pathway used DCNNs such as VGG-16 to yield stimulus encodings.^30^ They grouped convolutional layers of VGG-16 into 5 groups and found an homology between the outputs of groups and the BOLD responses along the early-to-late ventral visual stream.^30^ Inspired by this approach, we grouped the 13 convolutional layers into 5 stacks separated by max pooling layers. We then assessed the network’s performance on composite images with spatial attention applied to a stack of layers.

The trend of the multi-layer configuration is similar to that of the single-layer version: higher attention strength led to higher performance (Figure 4A, Figure 4—figure supplement 1A); attention could boost the network’s performance in response to low-contrast images (Figure 4B, left panel); and applying attention at later stacks produced greater performance increase (0.57 increase in F1 score at stack 5) (Figure 4C, Figure 4—figure supplement 1A). This is regardless of the modulation types (EAR, SUAR or EAR+SUAR). However, in contrast to the single-layer implementation, multi-layer EAR and SUAR could both effectively recover the model’s performance from signal degradation (low contrast) and noise addition (clutter) (Figure 4B, middle and right panels). Moreover, EAR exhibited comparable or even greater behavioral benefits than SUAR at every attention strength level (Figure 4A, B). In fact, EAR and EAR+SUAR attention outperformed SUAR at every stack with optimized attention strength (Figure 4C). On the contrary, when attention was applied to a single layer, EAR produced limited behavioral improvement and was outperformed by SUAR (Figure 2, Figure 3A). To further support such distinction, we selected the highest performance among single-layer implementations within a stack and compared it with the performance of modulating the entire stack. We found that EAR was always more beneficial when applied to multiple layers, whereas SUAR was not, except for stack 2 (Figure 4D). EAR+SUAR attention also produced higher performance when applied to a stack of layers except for stack 3. Additionally, EAR did not exhibit layer dependency when applied to a single layer (Figure 3A), but reached higher performance along the hierarchy when applied to a stack of layers (Figure 4C).

**Figure 4.**
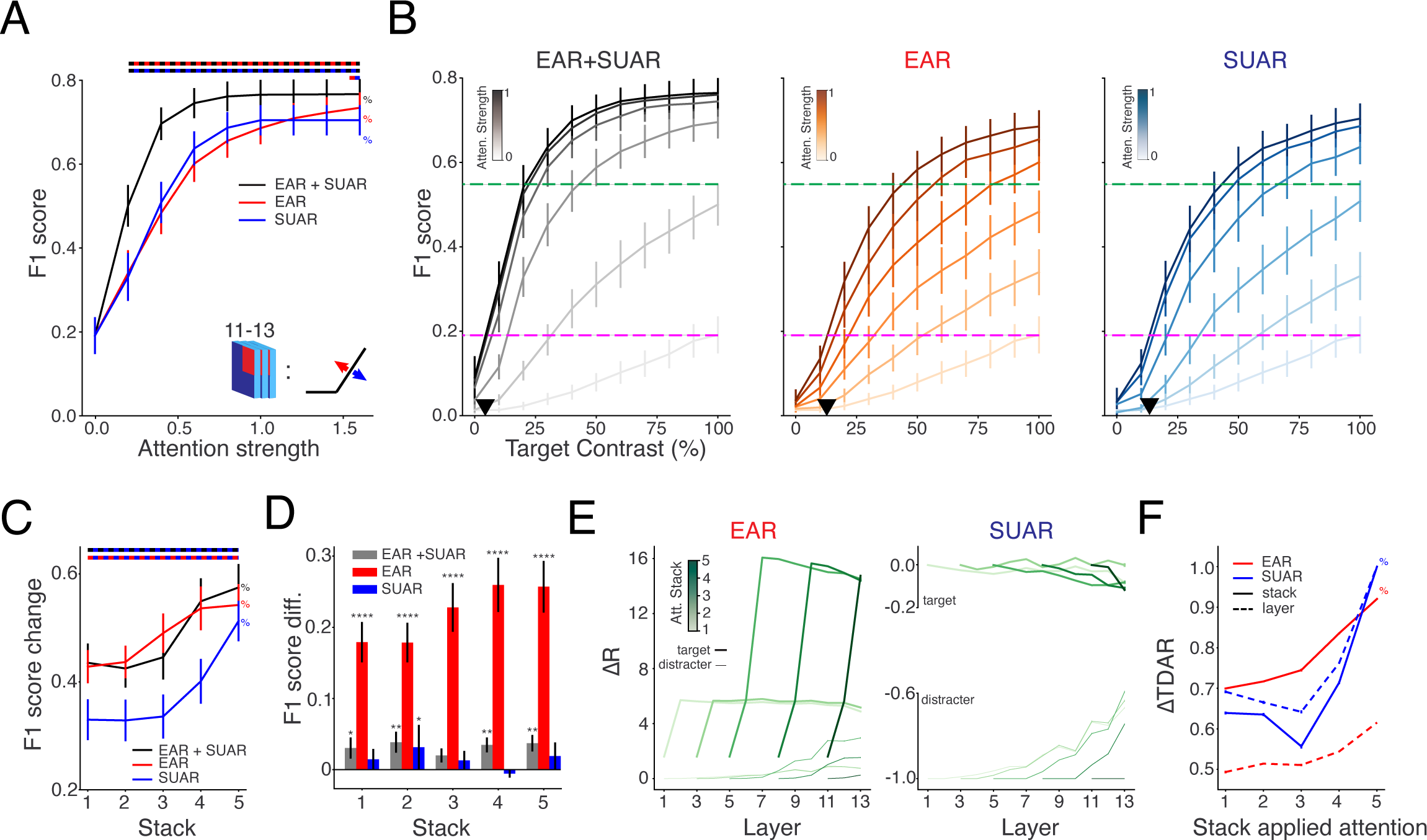
Effects of spatial attention applied to successive layers. (A) Mean network performance when attention of varying strengths was applied to the last stack of convolutional layers (layer 11-13). Attentional modulation was EAR+SUAR, or EAR, or SUAR. Horizontal bars on top indicate attention values for which there was a significant difference, with color code indicating the pairs of modulatory types considered for comparison (Wilcoxon signed-rank test, *N* = 20, Bonferroni corrected, *n* = 3, *p* < 0.0167). % indicates a significant difference in performance among different attention strengths for a given attention modulatory type (Friedman test with Conover post hoc pairwise comparisons, *N* = 20, Bonferroni corrected, *n* = 9, *p* < 0.00555 for any pair of comparison). (B) Mean network performance as a function of target contrast when applying different types of attention to the last stack of convolutional layers. Reference performances without attention are shown (dotted lines). Black triangle marks the lowest contrast level (maximal information loss) attention could recover, which is the contrast value of the intersection between the curve for highest attention and the magenta dotted line. (C) Average change in classification performance as a function of stack where attention was applied. For all modulatory types, best performing strength from the range tested (Figure 4—figure supplement 1A) was used for each stack. Horizontal bars on top indicate range of stack numbers for which there was a significant difference, with color code indicating the pairs of modulatory types considered for comparison (Wilcoxon signed-rank test, *N* = 20, Bonferroni corrected, *n* = 3, *p* < 0.0167). % indicates a significant difference in performance among different stacks for a given attention modulation type (Friedman test with Conover post hoc pairwise comparisons, *N* = 20, Bonferroni corrected, *n* = 5, *p* < 0.01 for any pair of comparison). (D) Difference in performance when an attentional type was applied to a stack vs. the best layer within that stack. The number of asterisks denotes the p-value cutoff of the significance of the performance difference for each modulatory type (Wilcoxon signed-rank test, *N* = 20), where * = *p* < 0.05, ** = *p* < 0.01, *** = *p* < 0.001, and **** = *p* < 0.0001. (E) Relative activity change (ΔR, Mean ± SEM, error bar is too small to be seen) in the target quadrant (thick lines) or distracter quadrants (thin lines) of each layer when EAR or SUAR was applied to stacks of layers. (F) Relative change of TDAR (ΔTDAR, Mean ± SEM, error bar is too small to be seen) of layer 13 when EAR (red) or SUAR (blue) was applied to different stacks (solid line). Dashed lines represent the largest ΔTDAR that can be achieved when attention was applied to individual layers within the corresponding stack. % indicates a significant difference in ΔTDAR among stacks (Friedman test with Conover post hoc pairwise comparisons, *N* = 20, Bonferroni corrected, *n* = 5, *p* < 0.01 for any pair of comparison). Significant differences were found between EAR-layer vs EAR-stack, SUAR-layer vs SUAR-stack, and EAR-stack vs SUAR stack for every stack, except for SUAR-layer vs SUAR-stack at stack 5 (Wilcoxon signed-rank test, *N* = 20, Bonferroni corrected, *n* = 4, *p* < 0.0125).

To understand the mechanism behind the improvement of multi-layer EAR, we analyzed unit activity change in each layer subjected to different attention conditions as we did for single-layer implementations (Figure 4E). We observed that the enhanced neural activity in the target region from EAR accumulated across layers: within the stack modulated by attention, the target-region activity change of the last layer was considerably higher than its previous layers (Figure 4E, left panel). Conversely, the extent of activity suppression from SUAR was bounded (−100%) so that the activity change in the distracter region remained the same for all layers within a stack (Figure 4E, right panel). In terms of the signal quality, both EAR and SUAR displayed robust layer dependency in ΔTDAR (Figure 4F). Because of the property of accumulation, EAR on multiple layers resulted in a much higher ΔTDAR compared to its effect on a single layer. When attention modulated stack 1-4, EAR boosted the TDAR significantly higher than SUAR, which agreed with their difference in the network’s performance (Figure 4C). Therefore, these analyses implied that EAR is more effective when applied to multiple convolutional layers because the gain of unit activity accrues along the hierarchy. On the other hand, SUAR does not benefit from the multi-layer implementation due to the existence of a floor effect, and it is most effective when applied to the last stage of the model. Note that at stack 1-4, multi-layer SUAR led to lower ΔTDAR than its single-layer counterpart (Figure 4F), which was probably due to the propagation of activity suppression from the distracter region to the target region. But such difference in ΔTDAR was not significant enough to drive a consistent difference in performance between the two versions of SUAR (Figure 4D).

## DISCUSSION

In this work, we use a DCNN model of the ventral visual stream to investigate the functional consequences of the modulatory effects of spatial attention across visual space and along the encoding stages. In the visual space with cluttered scenes, attentional modulation could be focal to the attended region or/and diffuse in the unattended region. Along the hierarchical encoding space, attention could focally modulate one processing stage or diffusely affect multiple areas. We find two alternative strategies that are equally effective at recovering information loss and reducing global noise: diffuse suppression of unattended region at the last stage of the hierarchical model, and focal enhancement of attended region across multiple encoding stages (Figure 5A). The behavioral benefits of attentional modulation correlate with the changes in unit activity and the resultant signal transmissibility in the DCNN, which provides a mechanistic explanation for the observed performance difference among attentional strategies. Specifically, unit activity enhancement accumulated along the hierarchy and led to a substantial increase in signal power. On the other hand, activity suppression could not build up beyond completely silencing the neurons. Thus, noise reduction was most effective at the last stage of visual processing.

**Figure 5.**
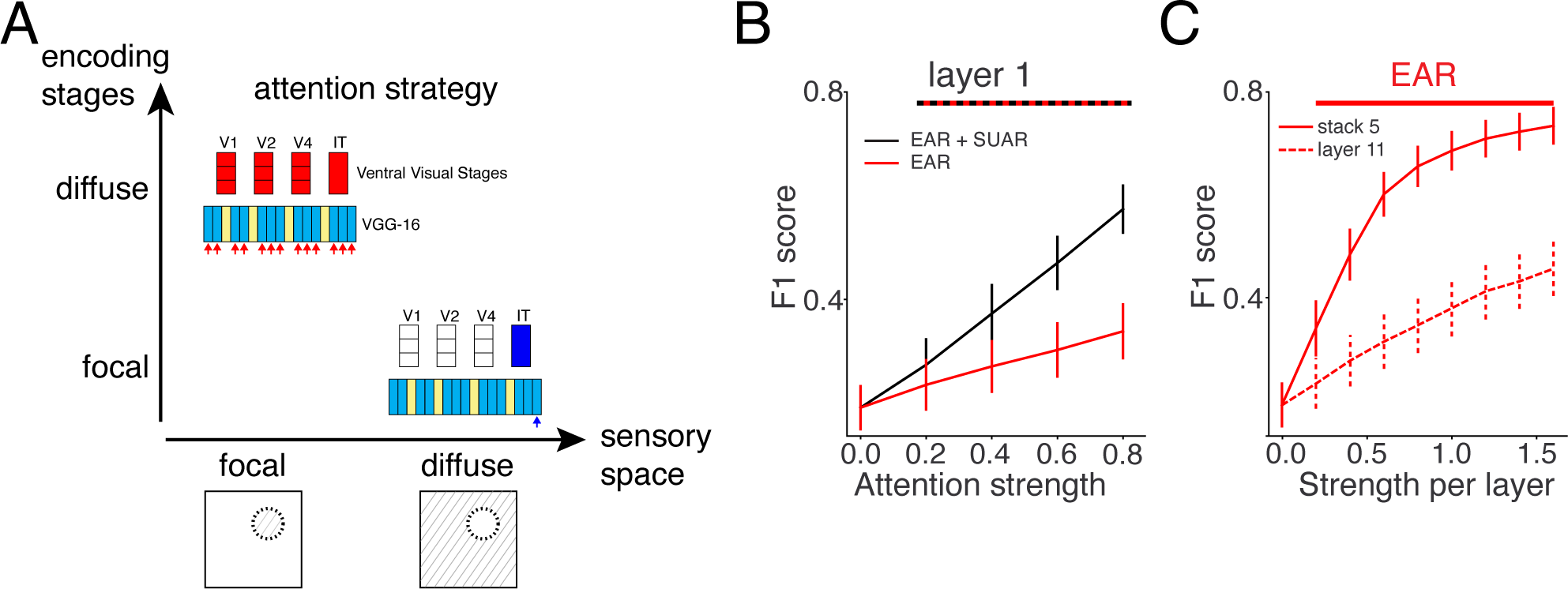
Focal vs diffuse strategies for attentional improvement in visual and cortical space. (A) Comparably effective strategies of attentional boosting of DCNN’s performance in a challenging task: diffuse suppression at the last convolutional layer (SUAR) or focal enhancement across multiple layers (EAR). The two strategies may correspond to SUAR in the higher visual region and EAR across the ventral visual hierarchy in the primate visual system. (B) Prediction 1: In an early visual processing stage, a combination of weak EAR and SUAR can achieve a significantly higher performance than EAR alone (significant attention values marked by the red-black line on top, Wilcoxon signed-rank test, *N* = 20, *p* < 0.05). (C) Prediction 2: With the same attention strength per layer, applying EAR to multiple layers consistently yields better performance than modulating a single layer (marked by the red line to top, Wilcoxon signed-rank test, *N* = 20, *p* < 0.05). A comparison example of stack 5 and layer 11 (first layer in stack 5) is shown.

Our results produce two key predictions for attentional modulation of neural activity observed in the ventral visual pathway. First, activity increase in a specific visual area, even though it accounts for only a small improvement in performance, could have a significantly greater impact if combined with unattended suppression (Figure 5B). Previous studies suggest that attentional increase of neural firing rate is modest in early visual areas^9, 16^ and could not account for a substantial fraction of the improved psychophysical performance.^2^ However, our results imply that the contribution of single-neuron modulations to behavior may be underestimated by considering rate increases in the attended region alone. Instead, the collective effects across visual space would be required to account for the full impact of attention. Attentional suppression of single-neuron activity has been found to exist at least in V1.^9^ Future recordings in other higher visual areas during attention tasks would be needed for testing our prediction about the functional roles of distractor suppression in performance improvement.

The second key prediction from our work relates to the multi-layer implementation of attention effects. Activity enhancement at multiple stages results in a much higher performance improvement than the same amount of modulation at a single stage (Figure 5C). Since attentional increase in firing rate has been reported in both striate and extrastriate cortices,^5, 16–18^ this result predicts that the net effect of activity enhancement on the signal-to-noise ratio of the ventral visual stream should be significantly larger than that estimated from a single processing stage.^2^ Consequently, the functional impact of rate modulation cannot be well appreciated by considering each brain region in isolation. Together, our results suggest that future investigation of attentional influence on neural activity in the primate visual system should encompass a relatively broader range of visual space that includes both attended and unattended regions and incorporate multi-area recordings across the hierarchy for a systematic analysis.

Our method of applying spatial attention was different from some other studies that implemented spatial attention either as a part of the network architecture or as a process of input selection,^54–59^ and whose aims were to obtain the state-of-the-art performance on benchmark tasks. The focus of our work was the distinct behavioral effects of attentional enhancement and suppression. Hence, the source of the attentional signal was not pertinent to our study and therefore not included in the model. Specifically, we modeled the attention effects by directly tuning the unit activity of the network instead of introducing top-down connections that exert attentional selection. Intriguingly, in a study that proposed an attention-based network for image caption generation, the attentional weights learned by the network concentrated on the object most relevant for the current word and ignored others,^58^ which was reminiscent of the target enhancement and distracter suppression that we implemented in this work. Future studies that examine the behavioral contribution of learned attentional weights in such networks would further support our thesis.

Mechanistically, we sought to understand the behavioral effects of attentional modulations by examining the relative contributions of signal and noise represented by the neural activity in the corresponding regions. It was the accumulation property that determined the advantage of the multi-layer implementation over the single-layer one for EAR but not for SUAR. Importantly, the capability of activity accumulation is built upon the non-saturating linear section of the activation function of the artificial neuron. However, a biological neuron has a physiological upper limit for its firing rate, defined by its absolute refractory period of action potential.^60^ But *in vivo*, biological neurons function in a dynamical range substantially lower than its physiological limitation.^61, 62^ Thus, attentional enhancement of neural activity in the brain is likely to accumulate along the visual pathway and our conclusion about the benefit of multi-layer EAR would also apply to the biological visual system.

We found that the most effective location to apply EAR+SUAR computations of spatial attention to the DCNN was the last stage of the processing hierarchy (Figure 3A, 4C), consistent with an earlier study on feature attention.^19^ These layers are thought to represent neuronal responses in visual areas V4 and LO^30^ that manifest strong response gain from attention.^49, 52, 63^ It is interesting to note that neuronal suppression from attention has been observed in the striate cortex,^7, 9, 11^ whose activity is better explained by the “early” stages of DCNNs.^30, 31^ Our study suggests that activity suppression decays with the depth of the network (Figure 3D). Therefore, we predict that attentional suppression also exists in higher visual areas to sustain its computational benefit. Future recordings in other visual stages are needed to test our prediction. Attention tasks that are appropriate for discerning suppressive effects should include a baseline attention condition, such as a broad distribution of attentional resources across the visual scene,^9^ so that neural activity change induced by directing attention away from the receptive field could be computed.

How attentional enhancement and suppression are implemented in the biophysical circuitry is still unclear. Since the frontal eye field area (FEF) is thought to be the source of spatial attention control signals (see Noudoost et al.^64^ for a review), feedback connections from FEF to posterior visual cortex as well as those from higher (e.g. V4) to lower (e.g. V1) visual areas are believed to achieve the neural modulatory effects of attention. Anatomical evidence suggests that these backward corticocortical projections target retinotopically matched areas.^65–67^ Therefore, we propose that attention fields share the same retinotopic location across the visual hierarchy and the enhancing effects at the attended region are cascaded across visual processing stages. Despite the numerous biophysical mechanisms for multiplicative response gain,^68–71^ the hierarchy of computations ensures the accumulation of response gain and its accompanying improvement of signal-to-noise ratio. As for the attentional suppression, the Selective Tuning model (ST), as a theoretical framework for explaining attention selection, posits that a suppressive surround in the unattended region arises as an inherent consequence of the top-down selection and pruning operation.^72–74^ In the visual processing hierarchy, a winner-take-all process selects the unit(s) that best represent(s) the attended input at each processing stage, and then prune(s) away other input connections from non-winning units to the winner. These pruned connections lead to the suppressive effect in the unattended region. ST predicts the dependence of attentional suppression on spatial scrutiny demand of the task (how narrow the focus of attention is required to be)^75^ and the existence of suppression even without distractors.^76^ But direct physiological evidence of the pruning operation is lacking. Mechanistically, one possible implementation is through the activation of local inhibitory interneurons, which are targeted by either the feedback connections directly^77, 78^ or the interlaminar projections.^79^ Another possibility by which the gain modulation of the neural response could be divisive is through the control of the background synaptic noise.^80, 81^ Glutamatergic inputs from feedback connections could target both pyramidal cells and local interneurons, generate balancing increase in background excitation and inhibition, and thereby reduce the evoked responses in the sensory area.

Another conclusion from our work relates to the dependence of attention effect on contextual information. Particularly, SUAR displayed more leverage for a more cluttered scene, whilst EAR produced context-independent effects (Figure 2—figure supplement 1D). Our finding contrasts with a recent study that characterized the relationships between the perceptual boost of attention and task-set properties.^82^ They showed that the level of visual clutter was not strongly connected to the perceptual boost of attention in the DCNN. However, their test images were original ImageNet images on which their model was trained, which was presumably the reason why their model was relatively insensitive to the clutter level in the image. Our task was different in that the composite images presented to the network introduced a new type of clutter that the network never encountered during training and thus substantially confounded the pre-trained network. Spatial attention to the network successfully compensated for the performance loss caused by such types of visual clutter, demonstrating the functional capability of attention in processing novel tasks.

Although in this study we assumed that a convolutional layer in VGG-16 corresponds to a visual area,^25^ the extent of biological neural circuit represented by a convolutional layer is still under debate. Beniaguev and colleagues demonstrated that a DCNN with 5-8 layers was required to fully capture the input/output function of a realistic cortical pyramidal cell.^83^ On a longer timescale, the output of a single convolutional layer or several sequential layers is able to predict the mean neural activity from a visual area in response to naturalistic images.^28, 30, 33^ Our findings about the difference between the multi-layer and single-layer versions of attention implementation provide another dimension to explore this question: the number of convolutional layers required to represent a visual area needs to exhibit the same activity-behavior relationship, i.e., a certain amount of neural manipulation (e.g., induced by attention) should induce the same difference in behavioral performance. Stimulating and silencing neurons in visual cortex separately could potentially offer insights into how attention-related neural enhancement and suppression influence the animal’s behavior.

Despite the high performance of object classification, the DCNN model we exploited lacks some neurobiological details such as recurrent and feedback pathways, cell types, and noisy and dynamical neural activity. There are four different levels of tasks with increasing difficulty that need to be addressed by any general-purpose visual system model: *detection, localization, recognition,* and *understanding*. Although a purely feedforward scheme of DCNNs matches or even surpasses human performance in object detection tasks, it is not sufficient to accurately localize objects because of the coarser representations of the output layer inherent in its pyramidal abstraction.^84^ The inclusion of feedback and recurrent connections in DCNNs shows that these additional information flows improve the model’s recognition capacity in more challenging situations,^34, 35, 38, 85^ increase the predictability for temporal dynamics,^35^ and develop visual features such as contrast adaptation and spatiotemporal representations.^86, 87^ Since feedforward connections establish the detection performance of DCNNs, we believe the main conclusion from our results, such as the functional distinction between enhancement and suppression, will generalize in more biologically-realistic models. We did not explore other attentional correlates of neural activity such as changes in trial-to-trial variability and noise correlations^2, 3^ since the input/output relationship of the artificial neurons in VGG-16 are described by a deterministic function. Further studies incorporating noisy spiking neuron models in a DCNN architecture^88^ will be helpful in elucidating the behavioral significance of modulating neural variability in artificial neural networks.

## METHODS

### Key Resources

The weights for the VGG-16 came from Frossard (2017)^89^ (RRID: <u>SCR_016494</u>). The weights for the binary classifiers came from Lindsay and Miller (2018)^19^ (Dryad: doi:10.5061/dryad.jc14081).

### Deep Convolutional Neural Network

The deep learning model we used to model the visual system is VGG-16, which is a convolutional neural network (CNN) that achieved 92.7% top-5 test accuracy in ImageNet 2014.^43^ VGG-16 is a feedforward architecture composed of stacks of convolutional and max pooling layers, followed by a sequence of fully connected layers (Figure 1A). The weights of the network were obtained by training on the standard ImageNet dataset including images from 1000 categories.^43^ The last fully connected layer contains 1000 units (one for each category) whose activities pass through a softmax classifier to generate the probability of the given image belonging to each of the 1000 categories.

The convolutional layer (as numbered in Figure 1A) contains a set of filters that convolves with the input from the previous layer to extract features. The unit activity of the convolutional layer is the rectified value of the convolution:

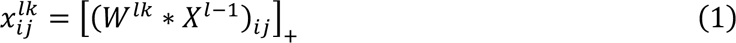

where [*x*]_+_ is the Rectified Linear Unit (ReLU) function, and * denotes convolution. *W^lk^* represents the weights of the *K^th^* convolutional filter in layer *l*. *X*^*l*-1^ is the output of all units from the previous layer 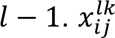 is the unit activity at the spatial location *i, j* from the *k^th^* feature map in layer *l*. Convolutional filters are 3 × 3, and the convolutional stride is fixed to 1 pixel. The spatial padding is set to ensure that the spatial resolution is preserved after each convolutional layer.

Max pooling layers slide a 2 × 2 filter over the outputs from the convolutional layer and select the maximum element from each feature map covered by the filter. They effectively reduce the dimensions of the feature maps for the following convolutional layers (layer 1 and 2: 224 × 224; layer 3 and 4: 112 × 112; layer 5, 6, 7: 56 × 56; layer 8, 9, 10: 28 × 28; layer 11, 12, 13: 14 × 14).

Two fully connected layers follow the last convolution layer and connect all units of the convolutional layer. The variant of VGG-16 we used in this study is different from the original model in that the third fully-connected layer of the original model is replaced with a series of binary classifiers, one for each of the 20 categories: paintbrush, wall clock, typewriter, paddlewheel, padlock, garden spider, long-horned beetle, cabbage butterfly, toaster, greenhouse, bakery, stone wall, artichoke, modem, football helmet, stage, mortar, consommé, dough, bathtub. Binary classifiers were logistic regressions trained on the ImageNet 2014 validation set.^19^ Each training batch contains 35 true positive images for each category and 35 true negative images from the remaining 19 categories.

### Test Images

The test images were samples of 20 categories from the ImageNet 2014 validation set but were different from those training images for binary classifiers. The color contrast of an image was controlled by blending the image with its solid grey image (<u>Python Imaging Library</u>): An image of 70% contrast was the result of 70% of the original image blended with 30% of the solid grey image. To create a more challenging task, we scaled down the size of these test images from 224 × 224 pixels to 112 × 112 pixels and arranged one target category image with three images from other categories on a two-by-two grid (composite images). The test image set for each category was composed of 150 composite images, half of which contained the target category (true positives) and the other half did not (true negatives). Both the precision and the recall of classifiers were calculated, and the performance was reported as the F1 score:

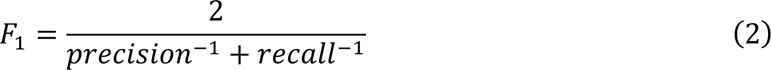

### Spatial Attention

In our study, we aimed to replicate the changes in neural activity by spatial attention on artificial units of a CNN to investigate their impacts on the model’s categorization performance. A convolutional layer in a CNN is designed as applying the same filter across the whole visual field to generate the feature map. Hence, the position of a unit within a feature map reflects the approximate center of its spatial receptive field. Even though the max pooling layer reduces the spatial dimensions of its following convolutional layers, the ‘retinotopy’ is still preserved throughout the network. Since the composite images for testing presented a target category image in a quadrant, we applied enhancing spatial attention effects to the layer units within the given quadrant (target category for “attended”) and suppressive effects to those in other three quadrants (nontarget categories for “unattended”).

Mathematically, attentional modulation of neural activity was through multiplication, corresponding to the response gain^49^ in non-human primate literature. We multiplied the slope of the rectified linear units (ReLU) by a strength parameter, β:

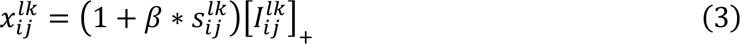

where 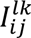 is the input from layer *l* − 1 and 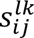 represents the direction of the modulation: 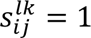 for units within the attended quadrant; 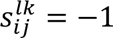 for units within unattended quadrants.

For a bidirectional attentional modulation, we applied both enhancement 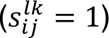 for the attended region and suppression 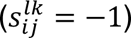 for unattended regions. Target enhancement (EAR) had no effect on unattended regions 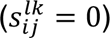 and distracter suppression (SUAR) kept neural activity in the attended region unchanged. Attentional modulation was implemented on an individual convolutional layer, a stack of layers, or all layers simultaneously.

### Analysis of layer activity

To understand the mechanism behind the improved performance by attentional modulation, we explored how a given activity change by attention propagates through the network by recording the neural activity and computing its relative change by attention of each convolutional layer in response to every test image:

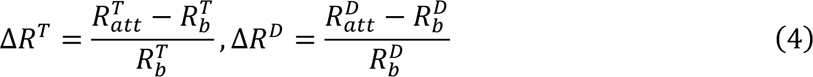

*R^T^* is the mean neural activity of the target region (quadrant for target image). *R^D^* is the mean neural activity of the distracter region (quadrants for distracter images). *R_b_* represents the mean activity without attention and *R_att_* is the activity after applying attention. For a given attention location, we used the attention strength that produced the highest performance. We averaged the activity change across spatial dimensions and images for every category and reported activity change as the mean across 20 categories.

To estimate the signal quality of the network, we assumed that the target signal was mostly encoded in the neural activity of the region that topologically corresponds to the target image, and the noisy signal was encoded by the neural activity of the distracter region. To reflect the signal-to-noise ratio, we computed the target-distracter activity ratio (TDAR), which is the negative ratio of the distracter region activity over the target region activity to avoid the appearance of 0 for the denominator:

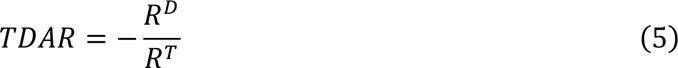

With this metric defined, we could derive the attention-induced relative change in TDAR from neural activity changes:

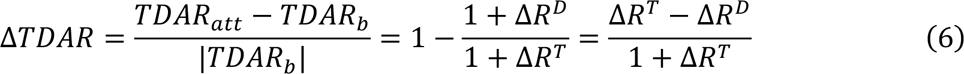

As shown in Figure 3, we focused on the TDAR of the last convolutional layer since it is the closest one to classifiers that possesses spatial information. TDAR robustly correlated the performance change induced by attentional enhancement and suppression, proving its validity as a measure of the signal quality of the network during information transmission.

---

**Figure 1—figure supplement 1.**
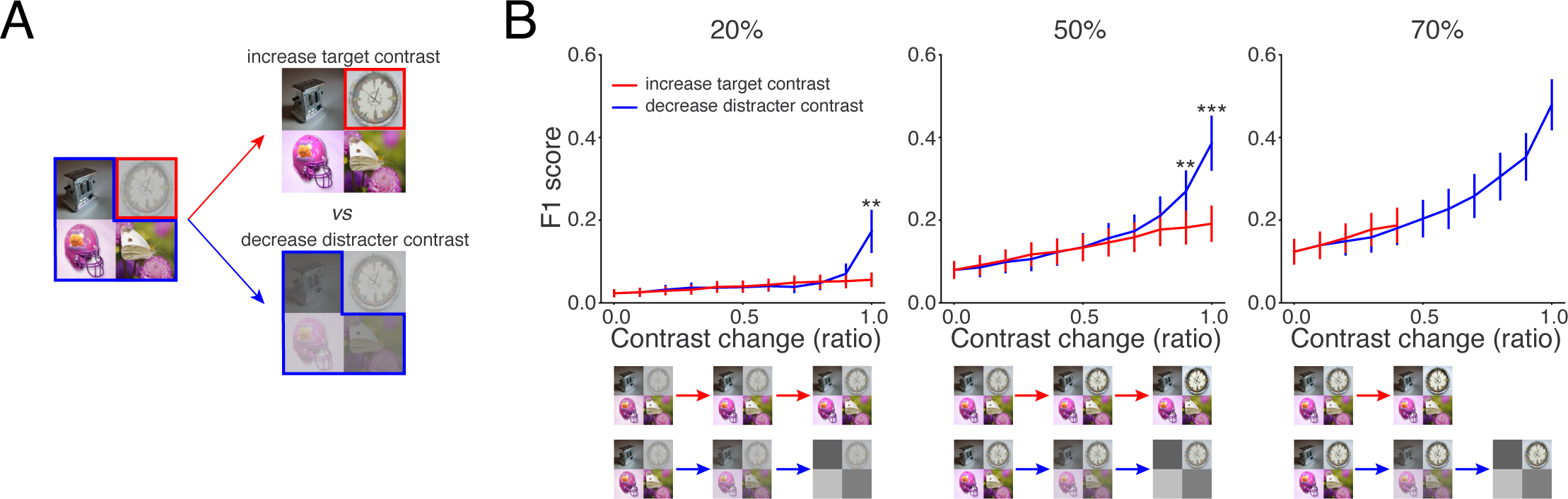
Effects of changing contrasts of the target or distracters on the network’s performance. (A) An experiment to probe the effect of tuning contrast in target or distracter images on the network’s performance. From a composite image with low-contrast target and full-contrast distracters, we either boosted the target contrast (red) or reduced the distracter contrast (blue) to see how it affects the performance. (B) The network’s performance on composite images with increased target contrast (red) or reduced distracter contrast (blue). We started from composite images with 20% (left), 50% (middle), or 70% (right) of target contrast. The contrast level was modified by a multiplicative ratio from 0 to 1. The reported performance was averaged across 20 categories (Mean ± SEM). The asterisk indicates a significant difference in F1 score between the two conditions (Wilcoxon signed-rank test, *N* = 20, * = *p* < 0.05, ** = *p* < 0.01, *** = *p* < 0.001, and **** = *p* < 0.0001).

**Figure 2—figure supplement 1.**
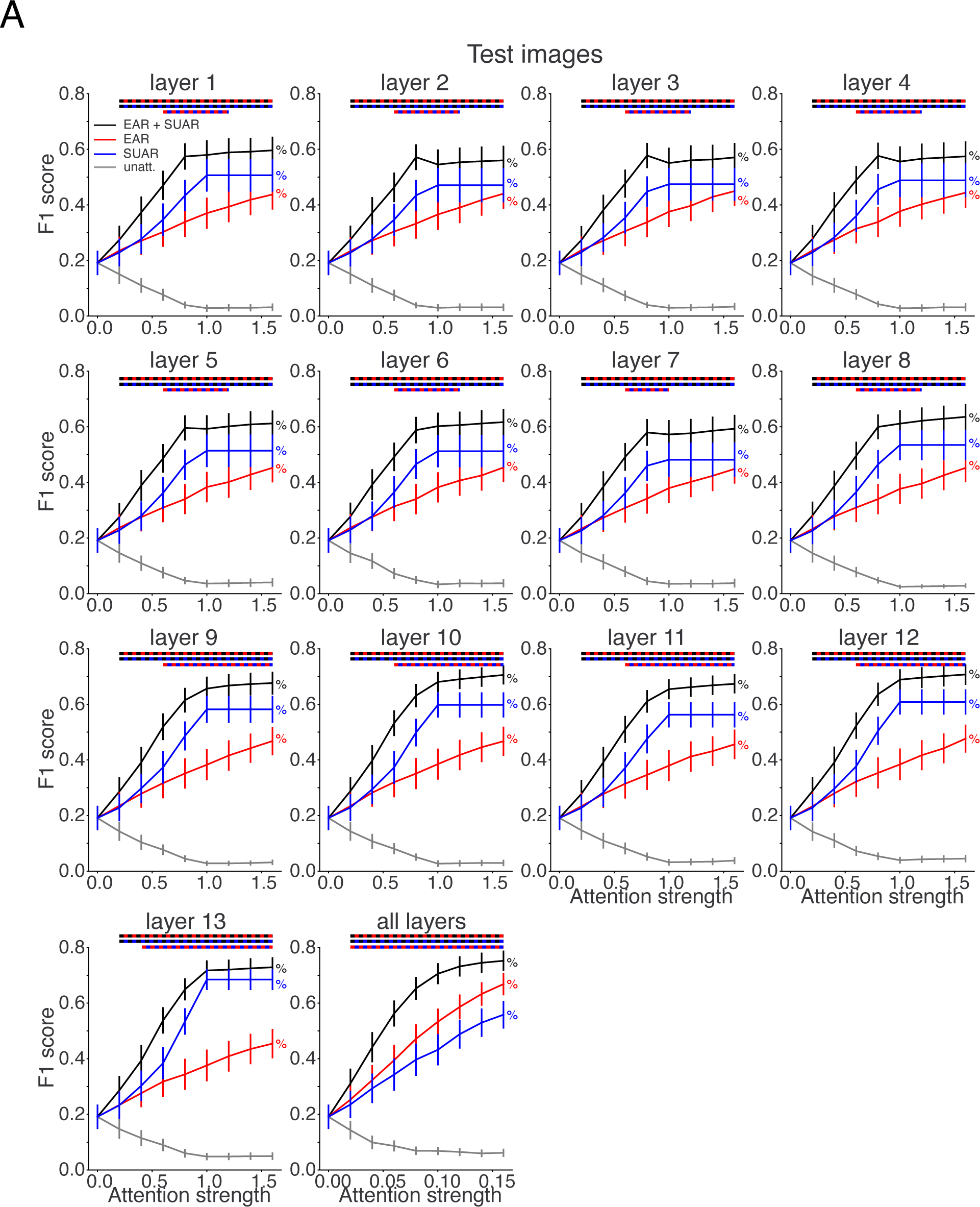
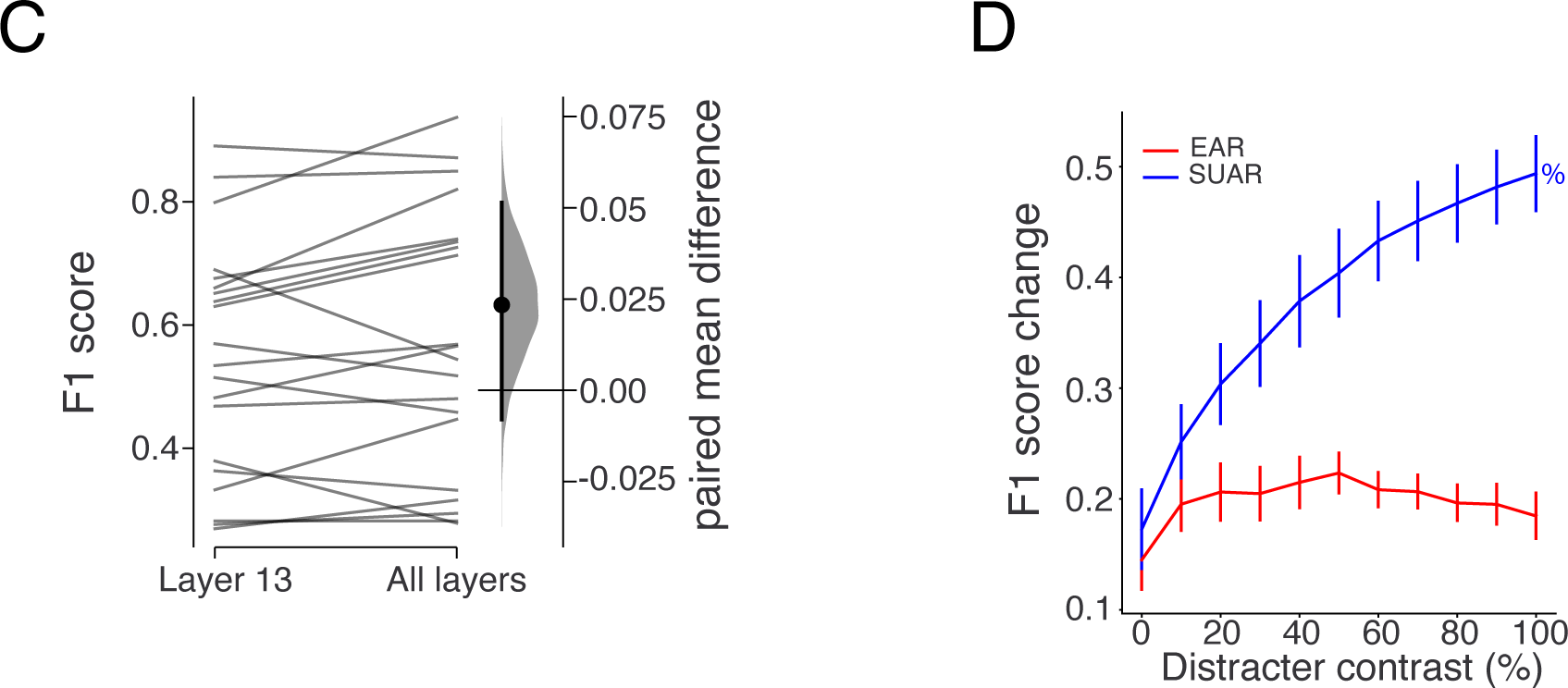
Attention effects at different locations or on different image sets. (A) Network performance as a function of attention strength when different types of attentional modulation (EAR+SUAR, EAR, SUAR, “unattended”) was applied to different locations. “Unattended” was a type of EAR+SUAR modulation when a distracter quadrant was “attended”. The reported performance was evaluated on full-contrast composite images and was averaged across 20 categories (Mean ± SEM). Dual-color dashed lines on top of each plot mark the range of attention strength that a significant difference between attentional types is detected (black-red: EAR+SUAR vs. EAR; black-blue: EAR+SUAR vs. SUAR; red-blue: EAR vs. SUAR; Wilcoxon signed-rank test, *N* = 20, Bonferroni corrected, *n* = 3, *p* < 0.0167). % indicates a significant difference in performance among different attention strengths for a specific attention condition (Friedman test with Conover post hoc pairwise comparisons, *N* = 20, Bonferroni corrected, *n* = 9, *p* < 0.00555 for any pair of comparison). (B) Same as (A) but on control images where distracter images have 0% contrast. (C) Gardner-Altman estimation plot showing the paired mean difference in performance when EAR+SUAR was applied to layer 13 vs. all layers. Both groups are plotted on the left axes as a slopegraph: each paired set of performance for a category is connected by a line. The paired mean difference is plotted on a floating axis on the right as a bootstrap sampling distribution. The mean difference is depicted as a dot; the 95% confidence interval is indicated by the ends of the vertical error bar. (D) Performance gain (Mean ± SEM) from EAR (red) or SUAR (blue) as a function of distracter contrast. Target contrast was 100% for all input images. Attention was implemented at layer 13 with the attention strength that optimized the performance for every contrast level. % denotes a significant performance difference across contrast levels for a specific attention condition (Friedman test with Conover post hoc pairwise comparisons, *N* = 20, Bonferroni corrected, *n* = 11, *p* < 0.00454).

**Figure 4—figure supplement 1.**
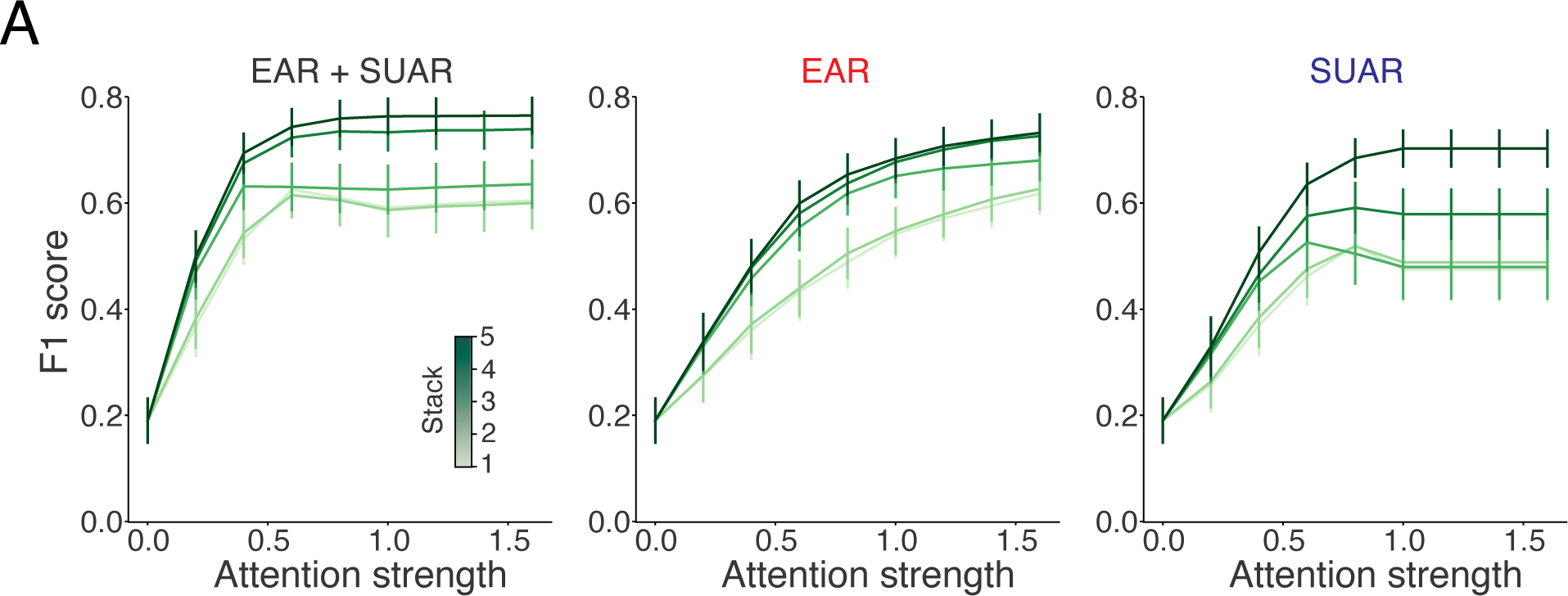
Multi-layer attention effects depend on attention strength. (A) Mean classification performance as a function of attention strength when spatial attention was applied to each of the 5 stacks of convolutional layers. Performance was evaluated for different types of attention and averaged across 20 categories (Mean ± SEM).

